# A novel Hsp90 phospho-switch modulates virulence in the major human fungal pathogen *Candida albicans*

**DOI:** 10.1101/2020.09.23.309831

**Authors:** Leenah Alaalm, Julia L. Crunden, Mark Butcher, Ulrike Obst, Ryann Whealy, Carolyn E. Williamson, Heath E. O’Brien, Christiane Schaffitzel, Gordon Ramage, Stephanie Diezmann

## Abstract

The ubiquitous molecular chaperone Hsp90 is a key regulator of cellular proteostasis and environmental stress responses. Hsp90 also regulates cellular morphogenesis, drug resistance, and virulence in human pathogenic fungi, which kill more than 1.6 million patients each year worldwide. Invasive fungal infections are difficult to treat due to the lack of effective antifungal therapies, resulting in mortality rates of up to 95%. As a key regulator of fungal virulence, Hsp90 is an attractive therapeutic target. However, fungal and animal homologs are highly conserved, impeding fungal-specific targeting. Thus, understanding the factors that regulate Hsp90 could provide an alternative strategy aimed at exclusively targeting this regulator of fungal virulence. Here, we demonstrate how CK2-mediated phosphorylation of two Hsp90 residues modulates virulence in a major fungal pathogen of humans, *Candida albicans*. We combined proteomics, molecular evolution and structural modelling with molecular biology to identify and characterize two Hsp90 phosphorylation sites. Phosphorylation negatively affects thermal stress response, morphogenesis, drug susceptibility and fungal virulence. Our results provide the first record of specific Hsp90 phosphorylation sites acting as modulators of fungal virulence. Post-translational modifications of Hsp90 could prove valuable in future exploitation as antifungal drug targets.

---

Fungi kill as many patients as tuberculosis and about three times as many as malaria each year^1^. One of the most deadly fungal pathogens, *Candida albicans* causes ~750,000 cases of invasive life-threatening bloodstream infections in immunocompromised patients worldwide each year^1^, with mortality rates approaching 75%^2^. In addition to high burdens on human life, fungal infections put a significant strain on health care costs. The United States alone spent $4.5 billion on 75,055 hospitalizations necessitated by fungal disease over ten years^3^ while English NHS trusts spend annually approximately £90 million on antifungal drugs^4^. This already dire situation is further exacerbated by the ever-growing number of patients at risk of contracting invasive fungal infections. It is thus imperative to understand the biological principles underpinning fungal virulence to identify points of fragility suitable for therapeutic targeting.

Diverse aspects of fungal pathogenicity are controlled by heat shock protein 90 (Hsp90), a ubiquitous regulator of the cellular protein homeostasis. Hsp90 regulates cell morphology, drug resistance, and virulence in *C. albicans*^5^ and other leading fungal pathogens of humans *Cryptococcus neoformans*^6^, which causes brain infections and *Aspergillus fumigatus*^7^ which targets the lungs, as well as the dermatophyte *Trichophyton rubrum*^8^, and *Sporothrix schenckii*^9^, the causative agent of cutaneous infections. Hsp90’s role in virulence and development is best understood in *C. albicans,* where pharmacological inhibition or genetic reduction of Hsp90 result in reduced drug resistance in planktonic cells^5^ and biofilms^10^, switching to filamentous growth^11^, induction of same-sex mating^12^, and delayed cell cycle progression^13^. Although attractive as a drug target for its role in fungal virulence, Hsp90 is highly conserved between mammals and fungi^14^. High sequence conservation prohibits specific targeting of fungal Hsp90 and resulted in severe host toxicity when mice infected with *C. albicans* where treated with Hsp90 inhibitors developed as anti-cancer therapeutics and antifungal drugs^15^. Thus, understanding fungal-specific regulatory mechanisms of Hsp90 could provide an alternative therapeutic approach and is consequently of biological and clinical relevance.

Post-translational modifications (PTMs) play a crucial role in regulating Hsp90. Their impact on chaperone activity, directionality of the chaperone cycle, and binding to co-chaperones have been extensively studied in mammalian cells and the model eukaryote *Saccharomyces cerevisiae*^16^. However, examples of Hsp90 PTMs in pathogenic fungi are limited to the observations that concurrent acetylation of two lysine residues, K30 and K271, resulted in increased susceptibility to Hsp90 inhibition, morphogenetic alterations, and impaired macrophage pyroptosis in *C. albicans*^17^, as well as attenuated virulence in *A. fumigatus*^18^. Given the importance of kinases as drug targets^19^ and experimental evidence of phosphorylation regulating most Hsp90 activities, this type of PTM is of particular interest. Phosphorylation negatively affects the chaperone cycle, interactions with clients and co-chaperones, conformational switching, and sensitivity to Hsp90 inhibitors^20^. The ubiquitous tetrameric casein kinase 2 (CK2), one of several kinases targeting Hsp90, phosphorylates Hsp90^T36^ in mammalian cells and Hsp90^T22^ in *S. cerevisiae*. This highly conserved residue was initially identified as critical for the survival of elevated temperatures^21^ and phosphorylation of T^22/36^ reduces kinase client stability and co-chaperone binding^22^. In *C. albicans*, CK2 phosphorylates Hsp90 serine and threonine residues^23^. Yet, specific Hsp90 residues targeted by CK2 and the functional consequences thereof have not been found.

Here we demonstrate that CK2-mediated phosphorylation of the newly identified Hsp90 residue S530 blocks *C. albicans* virulence and expression of virulence-related traits, such as cellular morphogenesis and survival of thermal stress. Probing the conserved T22 residue revealed that any alteration of this residue leads to loss of Hsp90 function. Our results emphasize the importance of Hsp90 phospho-regulation in fungal virulence and provide a potential site for future therapeutic targeting.

## Results

### Mapping, modeling, and evolutionary history of CK2 Hsp90 phosphorylation sites

To map CK2 phosphorylation sites in *C. albicans* Hsp90 *in situ*, the wild-type strain and four mutants, each lacking one CK2 subunit (for strain construction see Tables S1, S2, S3, and Supplementary Information), were grown in rich media at 30°C upon which epitope tagged Hsp90 was purified and analyzed by mass spectrometry. Without further enrichment, Hsp90 sequence coverage in trypsin digests was 70% or higher and ranged between 40-45% in endoproteinase Glu-C digests. Three phosphorylation sites were identified (Table 1). S279 was only phosphorylated in the *cka1*Δ/Δ mutant. S294 was phosphorylated in the wild type and all four CK2 mutants. S530 was phosphorylated in the wild type, the *cka1*Δ/Δ and the *ckb1*Δ/Δ mutants, but not the *cka2*Δ/Δ and *ckb2*Δ/Δ mutants. Interestingly, deletion of *CKA2* results in increased fluconazole resistance^24^, while Hsp90 inhibition abolishes fluconazole resistance^25^.

**Table 1:** Patterns of Hsp90 phosphorylation (P) detected in the wild type (WT) and the CK2 mutants.

| *C. albicans* genotypes |  |  |  |  |  |
| --- | --- | --- | --- | --- | --- |
| Hsp90 site | WT | *cka1Δ/Δ* | *cka2Δ/Δ* | *ckb1Δ/Δ* | *ckb2Δ/Δ* |
| S279 | - | P | - | - | - |
| S294 | P | P | P | P | P |
| S530 | P | P | - | P | - |

Thus, we hypothesized that phosphorylation of Hsp90^S530^ blocks Hsp90 function, thereby reducing resistance to antifungal drugs. The acquisition of a PTM that reduces drug resistance appears counter-intuitive, but the trade-off theory of optimal virulence predicts selective pressure to evolve reduced virulence in cases where the host and pathogen’s fitness are aligned^26^. *C. albicans’s* primary ecological niche is as a commensal on mucosal surfaces of the oral cavity and GI tract. To test this, we assessed the effect of Hsp90^S530^ phosphorylation on diverse virulence traits, including high temperature growth and morphogenesis.

In addition to investigating S530, we decided to also characterize the conserved CK2 target Hsp90^T22/T36^, whose *C. albicans* homolog is T25, called T25 henceforth. T25 resides in a highly conserved region of Hsp90’s N-terminus, which contains the ATP binding site and is the target of the Hsp90 inhibitors geldanamycin and radicicol^27^. S530 is located in a highly divergent region of the Hsp90 middle domain, where client proteins bind^27^ (Fig. 1a). To elucidate the relationship of these two residues within the Hsp90 molecule, we modeled the full-length *C. albicans* Hsp90 monomer on the *S. cerevisiae* crystal structure^28^ (Fig. 1b). A total of 609 residues (94% coverage) were aligned with 100% confidence, sharing 89% identity between both yeast species. The homology model revealed that both residues are located in loop regions without secondary structural elements (Fig. 1b). T25 resides in close proximity to the ATP binding pocket, while S530 is located in a loop near the protein’s C-terminus. The surface model shows that T25, although not a hydrophobic amino acid, is not solvent exposed, while S530 is (Fig. 1b). These observations are in keeping with molecular evolution theory, which predicts that decreased solvent accessibility results in fewer amino acid polymorphisms^29^ and thus higher sequence conservation.

**Figure 1:**
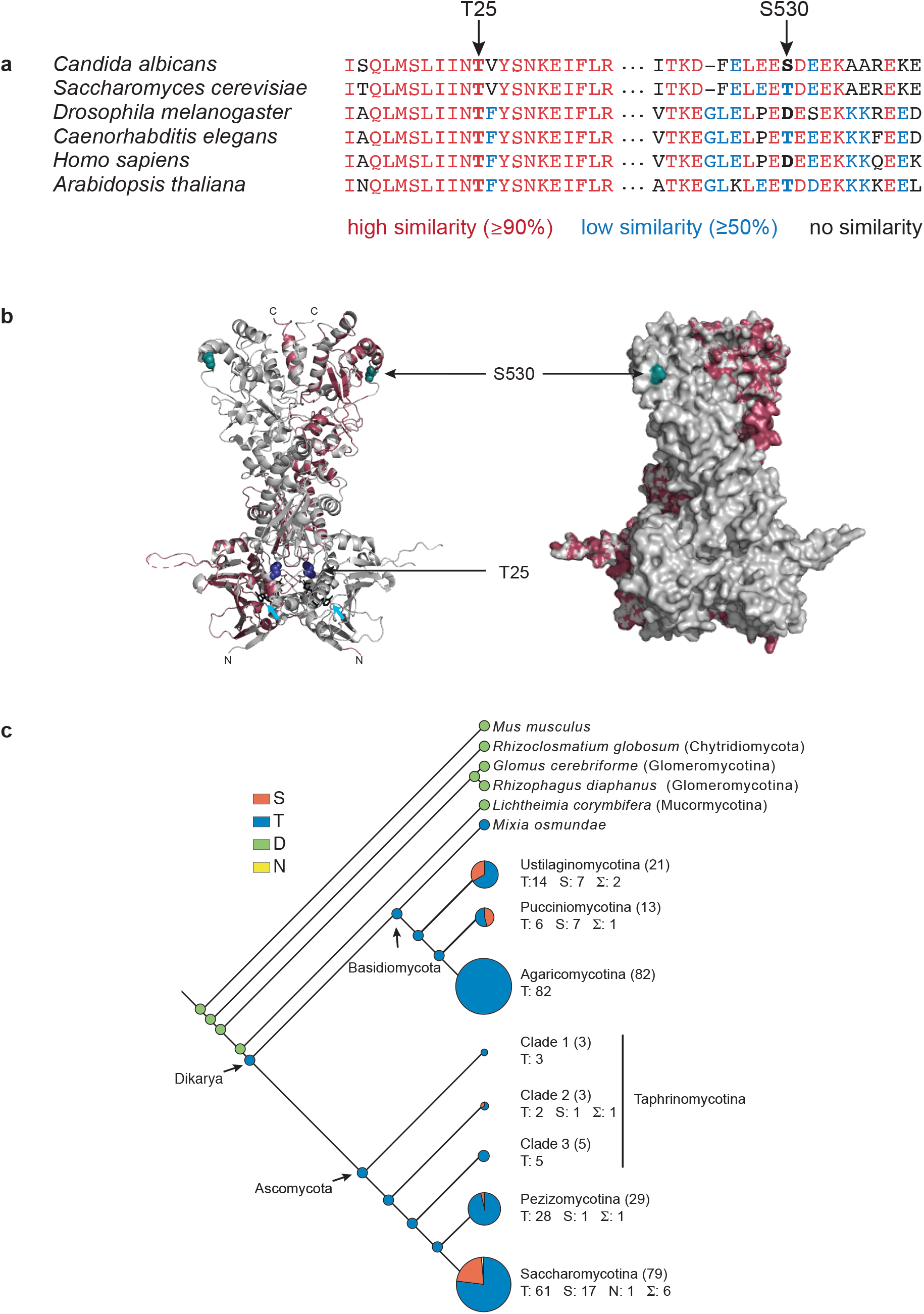
Structural modeling and ancestral character state reconstruction of CK2 phosphorylation sites in Hsp90. a) Alignment of eukaryotic Hsp90 protein sequences, comparing two fungal sequences to three animal and one plant protein. T25 resides in a highly conserved (red) region of the N-terminal domain of Hsp90. S530 is located in a divergent (blue) region of the Hsp90 middle domain. b) *C. albicans* Hsp90 (purple) modeled on the full-length crystal structure of *S. cerevisiae* Hsp90 (grey). T25 (blue) and S530 (teal) are visualized on the homology model (left) and the surface model (right). Azure arrows point to bound ATP in the homology model. c) Ancestral character state reconstruction of Hsp90^S530^ in 240 fungal taxa reveals at least eleven independent transitions from threonine (blue) to serine (red) within the Dikarya. Basal fungi share the ancestral aspartic acid residue (green) with the animal root (*Mus musculus*) of the tree. Within the Ascomycota and the Basidiomycota, subphyla have been collapsed and the circle size indicates the number of taxa within each clade. The number of taxa per subphylum is given in brackets, and the number of taxa with either threonine or serine or asparagine (yellow) are indicated with T, S, or N. The number of independent transitions within each subphylum is indicated by Σ. For character state reconstruction within ascomycetous and basidiomycetous subphyla see Fig. S1.

Given the divergent nature of the S530 residue (Fig. 1a), we investigated its evolutionary history within the fungal kingdom, asking if *C. albicans* presented a unique case or if multiple transitions could be detected. Aligning 659 amino acids of 240 fungal Hsp90 sequences, representing nine sub-phyla, with the murine homolog Hsp90ab1 revealed serine to be a derived allele originating at least eleven times independently within the Dikarya, which encompass the Ascomycota and Basidiomycota, and represent ~96% of known fungal species^30^ (Fig. 1c, S1, Table S4). Basal fungi, such as the Chytridiomycota, share an aspartic acid allele with animal cytosolic Hsp90. Threonine originated in the last common ancestor of the Dikarya and threonine to serine transitions were detected in five of the six sub-phyla. Serine alleles were detected in species considered plant pathogens (*Ustilago hordei*, *Puccinia* complex, *Aspergillus niger, Ashbya gossypii*), species associated with trees (*Neolecta irregularis*), fermentative yeast (*Nadsonia fulvescens*), methylotrophic yeasts *(Komagataella phaffi*, *K. pastoris*, *Ogataea parapolymorpha*, *O. polymorpha*), halo-tolerant species (*Debaryomyces hansenii, Zygosaccharomyces rouxii*) and the opportunistic pathogens (*Meyerozyma guilliermondii*, *Malassezia globosa*, *Candida parapsilosis*). No serine alleles were detected amongst the 82 Agaromycotina, which include the human pathogen *Cryptococcus neoformans* as well as mushrooms and jelly fungi. In summary, the serine allele is rare; it is present in less than 5% of fungi. Serine occurs in species that are capable of causing disease in plants, thrive in extreme environments, and exist as commensals of the human body where they can cause opportunistic infections in responses to changes in the host’s immune status, like *C. albicans*.

### Hsp90 phosphorylation negatively affects protein stability and co-chaperone binding

To investigate the effects of phosphorylation of T25 and S530 and to accommodate the ‘obligate diploid’ nature of *C. albicans*, we generated strains expressing one mutant *hsp90* allele, where the relevant residue was replaced with either a phosphomimetic glutamic acid (E) or a non-phosphorylatable alanine (A) while the remaining *HSP90* wild-type allele was placed under the control of the maltose-inducible promoter *MAL2*p (Tables S1, S2, S3, Supplementary Information). Consequently, strains grown in dextrose solely expressed the mutant allele, thereby revealing the effects of phosphorylation. Growth in maltose resulted in expression of the wild-type allele, serving as control and for strain maintenance. Non-phosphorylatable and phosphomimetic alleles were created for both residues (T25A, T25E, S530A, S530E) and compared to the wild type and the *MAL2-HSP90* promoter control, that carries two *HSP90* wild-type alleles, one of which is under the control of the *MAL2* promoter.

To test if phosphorylation affects Hsp90 stability, strains were grown in dextrose or maltose and cell extracts probed for Hsp90 using an antibody that recognizes amino acid residues 693-702 in the *C. albicans* Hsp90 C-terminus^31^. Replacing T25 or S530 with the phosphomimetic residue (E) almost completely depleted Hsp90 levels in cells grown in dextrose when compared to the *MAL2-HSP90* control. Protein levels remained comparable in the strains carrying the non-phosphorylatable alanine allele (A) (Fig. 2a). Thus, phosphorylation of T25 or S530 destabilizes Hsp90.

**Figure 2:**
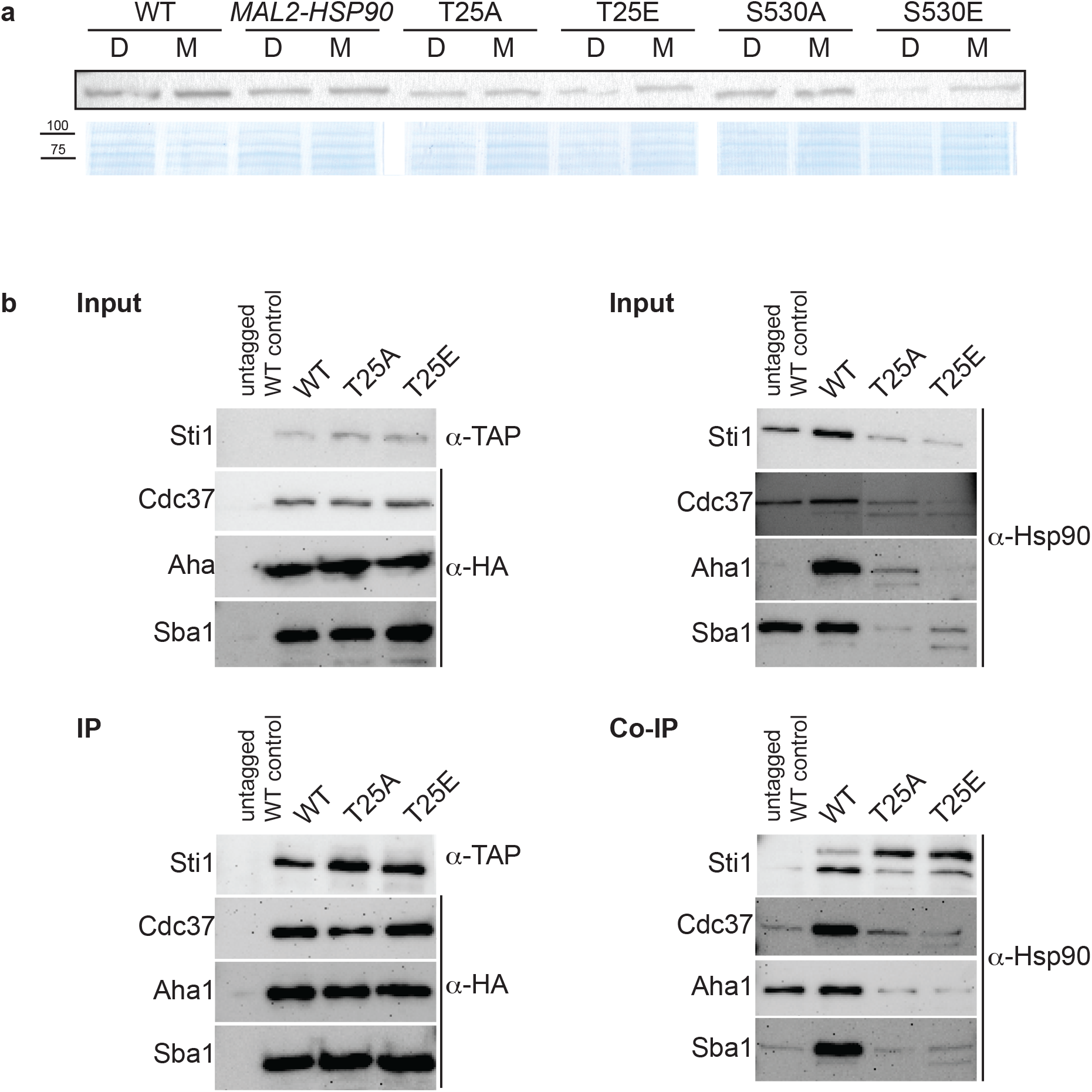
The *C. albicans* Hsp90 co-chaperone machinery is regulated by phosphorylation. a) One of two Western blots probed for presence of Hsp90 in *C. albicans* wild-type (WT), promoter control (*MAL2-HSP90*) and T25 and S530 mutant strains (top). Strains were grown in media containing either maltose (M) or dextrose (D) to regulate expression of mutant *HSP90* via the maltose-inducible promoter. A Coomassie-stained SDS gel validates equal loading of protein samples (bottom). b) TAP-tagged Sti1 and HA-tagged co-chaperones Cdc37, Aha1, and Sba1 were co-immunoprecipitated with Hsp90 in the wild type (WT) and the T25 mutants. The input was first probed for presence of co-chaperones and Hsp90 prior to immuno-precipitation with IgG or α-HA agarose targeting the co-chaperones. Levels of co-chaperones were then assessed in the immuno-precipitate (IP) using α-TAP and α-HA antibodies. To screen for co-immunoprecipitated Hsp90 (Co-IP), the membrane was probed with an Hsp90-specific antibody.

Given the importance of T22/T36 for co-chaperone binding in *S. cerevisiae* and mammalian cells^32^, we aimed to determine if the homologous T25 residue plays a similar role in *C. albicans*. Cell lysates from strains with TAP or HA tagged co-chaperones were immunoprecipitated with IgG or α-HA agarose and probed for presence of co-chaperones Cdc37, Aha1, Sba1, and Sti1, which bind to Hsp90’s N-terminal domain^33^, and for co-immunoprecipitated Hsp90 (Fig. 2b). Changing T25 to either a phosphomimetic or non-phosphorylatable form severely reduced binding of Hsp90 to all four co-chaperones, indicating functional conservation of T25 with regards to co-chaperone binding.

### Hsp90 phosphorylation blocks expression of key virulence traits

Mutational analysis of the *S. cerevisiae* homolog *HSP82* revealed T22 to be essential for survival of elevated temperatures^34^, a key virulence trait. To determine the importance of T25 and S530 for growth at high temperatures in *C. albicans*, *hsp90* mutants as well as the wild-type and promoter control strains were spotted onto medium containing either dextrose or maltose and exposed to different temperatures (Fig. 3a). Although neither T25 mutant grew as robust as the wild type or the *MAL2-HSP90* control, the phosphomimetic T25E grew better than the non-phosphorylatable T25A at temperatures above 25°C. Single colonies were detectable for T25E at the highest dilutions even at 37°C and 39°C. Growth of T25A was only detectable at the lowest dilutions at elevated temperatures. In either strain, however, colonies were visibly smaller than in the control suggesting reduced growth rate due to alterations of the T25 residue. Growth of the non-phosphorylatable S530A mutant was comparable to the control strains but the phosphomimetic S530E mutant grew much less robust especially at lower temperatures. This suggests phosphorylation of T25 and S530 plays different roles during exposure to thermal stress. Phosphorylation of T25 is required for survival of high temperatures, while phosphorylation of S530 impairs the cell’s ability to cope with lower temperatures.

**Figure 3:**
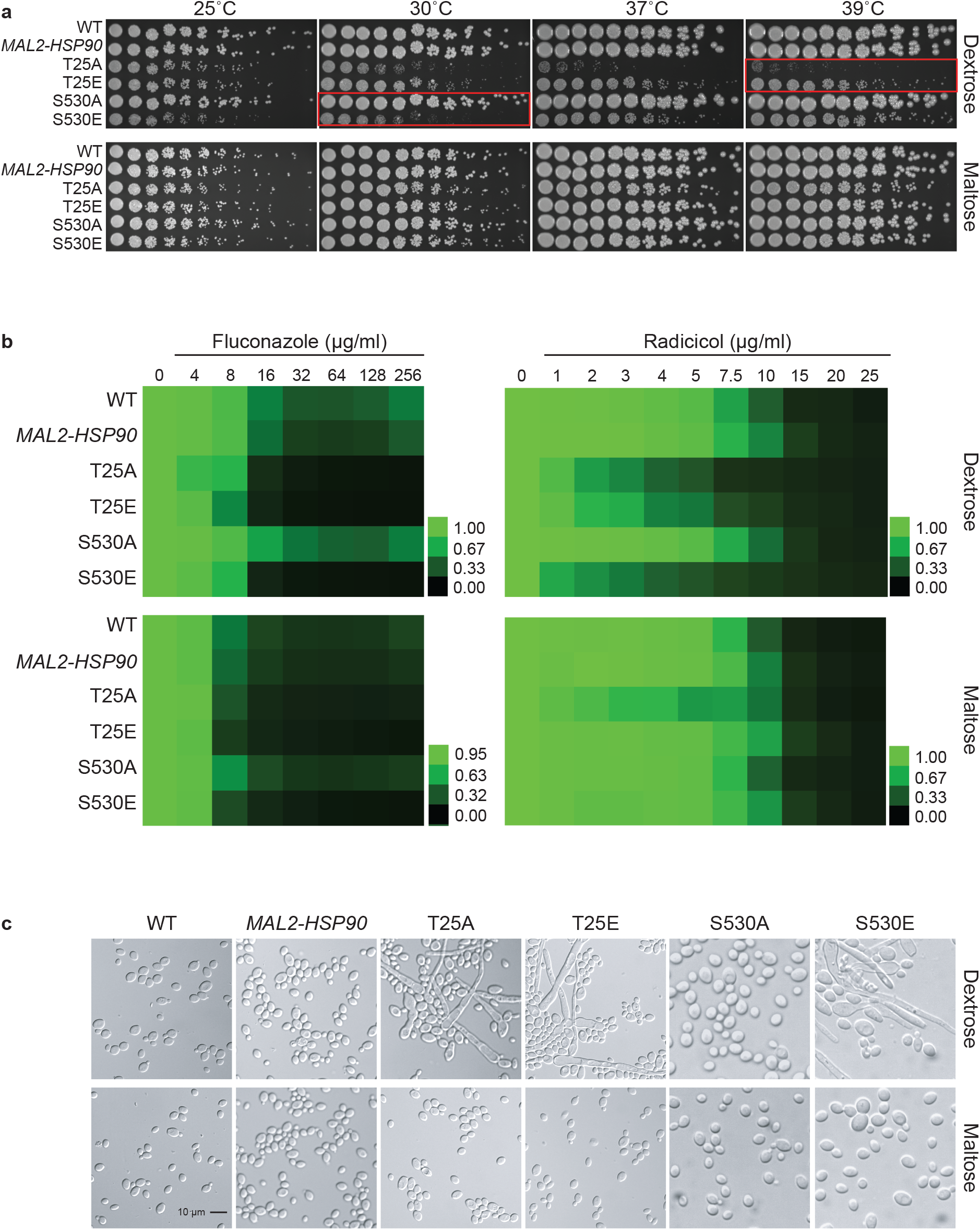
Expression of key virulence traits is contingent on Hsp90’s phosphorylation status. a) Two-fold serial dilutions of the *HSP90* wild-type, the *MAL2-HSP90* promoter control strain, and *hsp90* phospho-mutants were spotted onto solid media containing either dextrose or maltose and incubated at the indicated temperatures for 48 hours. Red boxes highlighting severe phenotypes. b) Wild-type and mutant cells were exposed to increasing doses of the commonly employed antifungal drug fluconazole and the Hsp90 inhibitor radicicol in media containing either dextrose (top) or maltose (bottom). Following incubation at 25°C, cell growth was assessed, normalized to the media only control and expressed as heat map where green indicates full growth and black represents no growth. c) DIC microscopic images of representative cells from each strain confirming morphogenetic changes of cells grown in dextrose (top) but not maltose (bottom). Shown are one of two biological replicates.

Given the importance of Hsp90 in *C. albicans* antifungal drug resistance and the role of Hsp90 PTMs in susceptibility to pharmacological inhibition of Hsp90 function, we tested the phospho-mutants for susceptibility to the commonly used antifungal drug fluconazole and the Hsp90 inhibitor radicicol. Control and phospho-mutant strains were exposed to drug gradients ranging from either 0-256 μg/ml for fluconazole or 0-25 μg/ml for radicicol. Growth was quantified after 48 hours at 25°C as optical density revealing that expression of *hsp90* mutant alleles sensitizes *C. albicans* to antifungal drug treatment and Hsp90 inhibition (Fig. 3b). Changing T25 to either a phosphomimetic or non-phosphorylatable allele or expressing the S530 phospho-allele rendered *C. albicans* susceptible to either drug. This suggests Hsp90 phosphorylation to be a determinant of drug susceptibility.

Cellular morphogenetic diversity is a fundamental element of *C. albicans’* virulence repertoire as the ability to switch between yeast and hyphae is essential for the infectious process^35^. To determine how phosphorylation of T25 or S530 affects cellular morphogenesis, the wild type, the *MAL2-HSP90* control and the *hsp90* mutants were grown in standard non-filament inducing conditions in rich media at 30°C. Microscopic examination of planktonic cells showed robust hyphal growth in both T25 mutants as well as the S530E strain in media containing dextrose, but not maltose (Fig. 3c). To quantify these observations, we calculated the morphology index (MI) for 100 randomly selected cells for each strain (Fig. S2). An MI > 1 is indicative of cell elongation or hyphal growth. MI scores for cells grown in dextrose or maltose differed significantly for both T25 mutants and S530E. In maltose, MIs clustered below 1, in dextrose MIs ranged from ~0.2 to ~75. Hence, both T25 mutant alleles and the phosphomimetic S530E allele cause cells to switch from yeast to hyphal growth.

### Hsp90’s phosphorylation status affects colony appearance and biofilm viability

Following up on the observation that differential Hsp90 phosphorylation results in elongated cell shape, we assessed macroscopic changes to colony morphology and response to cell wall stress. To capture changes in colony morphology, all strains were grown on nutrient-limited RPMI, synthetic defined (SD) media alone and supplemented with Calcofluor White or Congo Red as well as Spider media, which induces the yeast-to-hyphae transition (Table S5). Upon visual inspection of plates grown for 48 hours at 30°C, wild-type colonies were indistinguishable from the *MAL2-HSP90* control (Fig. 4). Colonies for both T25 mutants and the S530E strain appeared different from controls on RPMI and SD media. Despite being highly filamentous on a cellular level, colonies of these strains lost their hyphal edge on RPMI. On SD, these colonies were wrinkly rather than smooth. Growth on maltose restored wild type-like growth on RPMI and, to a degree, on SD media. No appreciable differences were observed for colonies grown in the presence of Congo Red or Calcofluor White when compared to SD only media (data not shown). Differences in colony morphology on Spider media appeared to be driven by the carbon source rather than a strain’s *HSP90* allele.

**Figure 4:**
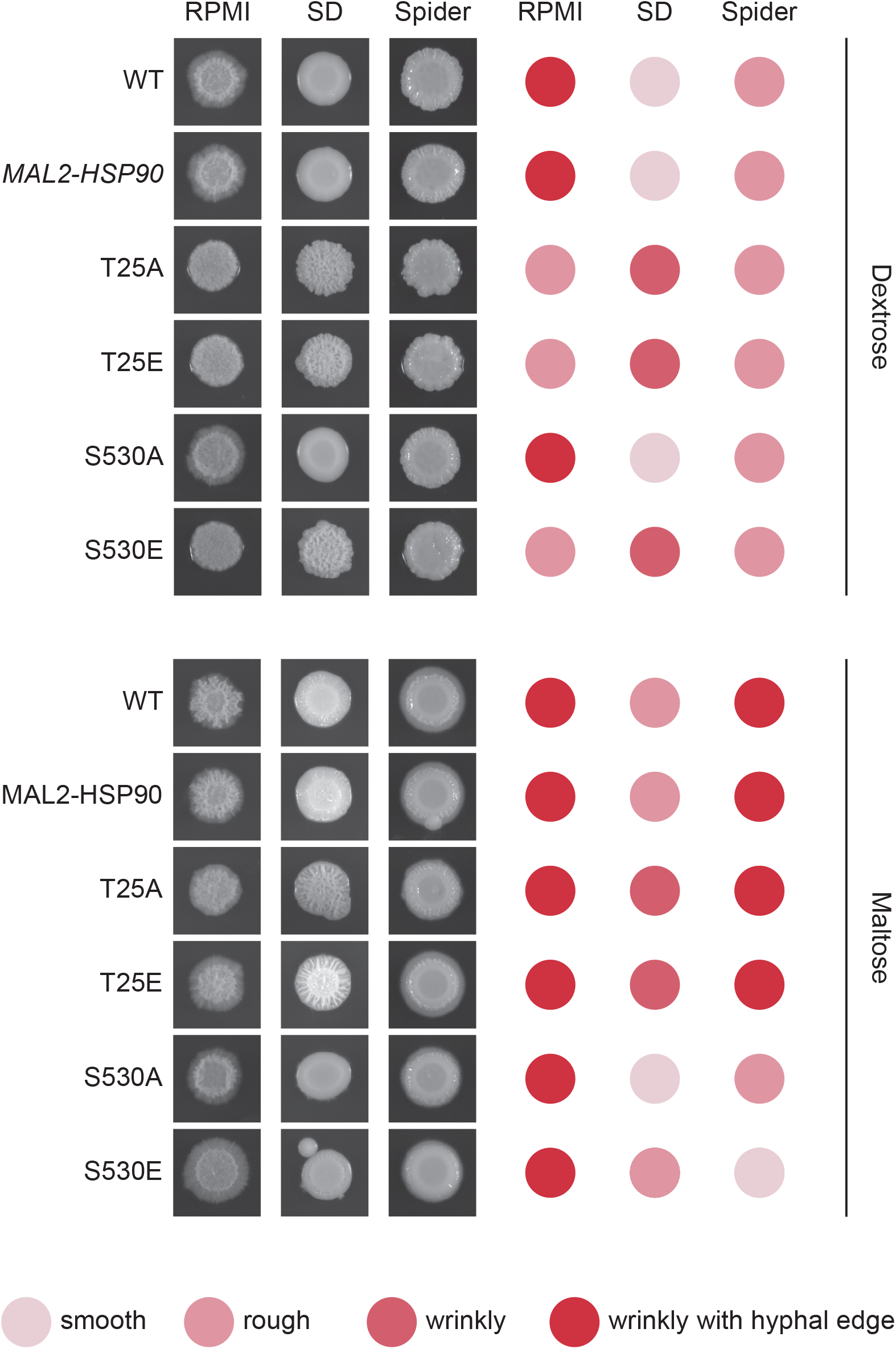
Aberrant colony morphology in response to changes in Hsp90 phosphorylation patterns. Images of colonies spotted onto RPMI, Spider or synthetic defined (SD) media containing either dextrose (top) or maltose (bottom). Colonies were scored for their appearance. The darker red the dot, the more extreme the colony phenotype. Shown is one of two replicates.

Morphogenetic diversity is a key prerequisite for the formation of *C. albicans* biofilms, which are medically relevant for two reasons. First, biofilms act as reservoirs that continuously feed the infection, and secondly they are intrinsically resistant to treatment with antifungal drugs^36^. To determine if Hsp90’s phosphorylation status affects the intricate developmental program underlying biofilm formation, cells were grown in standard biofilm-inducing conditions in RPMI (2 g/l glucose) at 37°C. Mature biofilms were imaged using scanning electron microscopy and their cell viability and biomass were quantified employing spectrometric approaches. Microscopic analysis of biofilm architecture at 800x and 8,000x magnifications showed that control strains and phospho-mutants form structurally similar biofilms, suggesting that Hsp90’s phosphorylation status does not fundamentally affect overall biofilm development in the conditions tested (Fig. 5a). More detailed inspection of biomass production and cell viability revealed that the latter is significantly decreased in the S530E mutant (p=0.00205) relative to the *MAL2-HSP90* control (Fig. 5b, c). This is in keeping with previous observations of phosphorylation of Hsp90^S530^ resulting in a loss-of-function phenotype. All other phospho-mutants displayed significant increases in cell viability (T25A: p=4.01×10^−14^; T25E: p=9.94×10^−13^; S530A: p=8.68×10^−7^). While deviating from our observation of a growth defect in the T25 mutants at 37°C on YPD agar, Azadmanesh *et al*. have shown that the physical environment influences *C. albicans* gene expression and development^37^.

**Figure 5:**
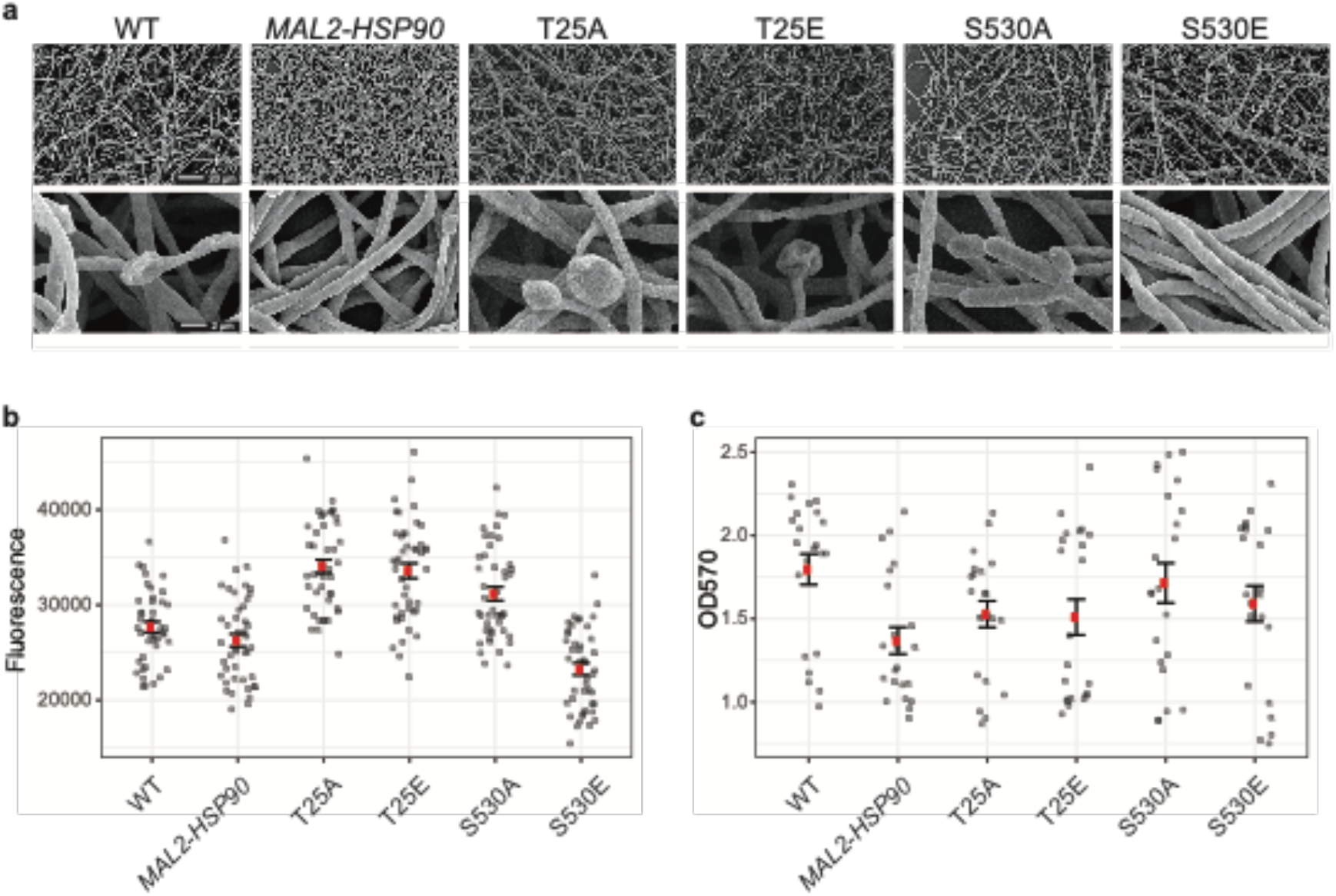
Biofilm architecture remains unaffected by Hsp90 phosphorylation. a) Scanning electron microscopy images of mature biofilms at 800x (top) and 8,000x (bottom) magnification reveal biofilm architecture between the wild type, the control and the phospho-mutants to be indistinguishable upon visual inspection. b) Biofilm cell viability, measured as fluorescence for 44 replicates per strain, is significantly reduced in strain S530E. c) Metabolic biomass measurements for 22 replicates per strain as absorption at OD570 were comparable to the promoter control.

Building on our finding that Hsp90’s phosphorylation status determines its effect on *C. albicans* cellular morphogenesis, we demonstrated that phosphorylation of S530 results in changes in colony morphology and reduced biofilm cell viability due to loss of Hsp90 function. Any manipulation of the T25 residue, however, leads to elongated cells, aberrant colony morphology and increased biofilm cell viability.

### S530 regulates *C. albicans* virulence in an invertebrate model of fungal pathogenesis

Once we established the importance of phosphorylation of S530 and T25 in virulence-relevant traits using different *in vitro* assays, we aimed to determine if and how Hsp90 phosphorylation affects fungal virulence. To test this, *Manduca sexta* caterpillars, an invertebrate model host of fungal disease^38^, were infected with 10^7^ yeast cells. Caterpillar survival was then assessed in groups of ten after 16 hours of incubation at the physiologically relevant temperature of 37°C. Strain S530E displayed significantly reduced virulence compared to the *MAL2-HSP90* control (Table S6). Nine caterpillars infected with S530E survived compared to two caterpillars infected with *MAL2-HSP90* (p=0.0054775). Thus, S530 phosphorylation blocks Hsp90 function, thereby attenuating virulence in an animal host.

In summary, Hsp90’s phosphorylation status affects fungal virulence. Taken together, our results suggest that S530 is a novel phospho-switch. When phosphorylated, it blocks Hsp90 function and thus the expression of virulence traits, which may aid its adaptation to a commensal lifestyle while maintaining the ability to cause invasive infections. Alterations of T25, however, are important beyond phosphorylation, suggesting this residue to be a ‘switch point’, important for structural integrity of *C. albicans* Hsp90.

## Discussion

Our data indicate that CK2-mediated phosphorylation of Hsp90^S530^ blocks expression of virulence traits in a major human pathogen while any alteration of Hsp90^T25^ resulted in a loss-of-function phenotype. This suggests that S530 is a novel *C. albicans* Hsp90 phospho-switch while T25 acts as switch point.

Similarly, to what has been observed in *S. cerevisiae* and mammalian cells, any modification of *C. albicans* T25 negatively affects co-chaperone binding, survival of high temperatures, and susceptibility to drug and Hsp90 inhibition^32,39^. We thus propose that T25 acts as a switch point in *C. albicans* Hsp90. Switch points impact on the overall conformation and function of a protein through conformational disturbances altering chaperone function^40^. These types of allosteric PTMs are of evolutionary significance as they expand the proteome’s complexity and facilitate dynamic responses to a large number of stimuli without the need for additional genes^41^. The effect, however, appears to be restricted to T22 in *S. cerevisiae* and T25 in *C. albicans*, as changes in the acetylation status of the neighboring conserved *C. albicans* K30^17^ or *S. cerevisiae* K27^42^ residues did not result in loss of Hsp90 function.

Detection of phosphorylation of S530 has remained rather elusive. Only one^43^ of five global phospho-proteomic analyses of *C. albicans*^44–47^ detected phosphorylation of Hsp90^S530^ in *C. albicans* hyphal cells^43^. This finding is consistent with our experimental data showing filamentous growth in the phosphomimetic S530E mutant. While our proteomic analysis revealed S530 to be phosphorylated in the wild type, the wild-type and *MAL2-HSP90* control are phenotypically more similar non-phosphorylatable mutant S530A than the they are to phosphomimetic mutant S530E (Fig. 3). This would suggest that phosphorylation of S530 is sparsely deployed and dynamic. Characterizing the *S. cerevisiae* homologous residue T533 using phosphomimetic and non-phosphorylatable alleles revealed that this residue does not affect survival at elevated temperatures, susceptibility to Hsp90 inhibition, or yeast growth^48^ suggesting evolutionary divergence in sequence and function. Taken together, the dynamic phosphorylation status of S530 carefully orchestrating *C. albicans* virulence in a commensal turned pathogen yeast and the lack of involvement in Hsp90 function of T533 in the benign yeast *S. cerevisiae* would be in keeping with S530 to be a genetic factor that reduces severity as predicted by the trade-off hypothesis of optimal virulence theory.

In conclusion, Hsp90’s phosphorylation status and structural integrity affect fungal virulence. CK2-mediated phosphorylation of the divergent S530 residue resulting in reduced fungal virulence could be further exploited as a novel therapeutic target to fight fungal infections.

## Methods

### Strains, strain construction and culture conditions

All strains used in this study are listed in Table S1. A detailed description of strain and plasmid construction can be found in the Supplementary Information together with Tables S2 and S3, which list the necessary primers. In preparation for experiments, strains were grown for 16-18 hours in YPM (1% yeast extract, 2% peptone, 2% maltose) at 30°C while shaking at 200 rpm, unless specified otherwise. To repress the *HSP90* wild-type allele, strains were grown in YPD (1% yeast extract, 2% peptone, 2% dextrose) at the temperatures specified. For long-term storage, strains were maintained at −80°C in 25% glycerol.

### Identification of the S530 phosphorylation site

The S530 phosphorylation site in Hsp90 was identified by mass spectrometric analysis of purified Hsp90 in the Proteomics Core Facility at EMBL Heidelberg, Germany. To purify Hsp90 from CK2 deletion mutants (YSD692, YSD675, YSD694, YSD696) and the corresponding wild type (YSD673), strains were grown to mid-log phase at 30°C in 200 ml volumes in YPD at 30°C. Whole cell proteins were extracted as described below and the C-terminally TAP-tagged chaperone was pulled down using IgG agarose beads from lysate containing 20 mg of protein (see below). The protein was eluted in 100 μl 1x sample buffer and heated to 95°C for 10 minutes prior to loading 30 μl onto a 10% Tris-glycine SDS-PAGE gel for separation. Coomassie-stained gels were imaged and the bands corresponding to Hsp90 were cut out and processed by in-gel digestion^*49*^. Hsp90’s phosphorylation sites were identified using Glu-C and trypsin digestion and liquid chromatography-mass spectrometry on a QExactive plus mass spectrometer (Thermo Fisher).

### Structural modelling

A model of the full-length *C. albicans* Hsp90 monomer was generated based on the *S. cerevisiae* Hsp90-Sba1 crystal structure^28^ (PDB #2cg9) using the Phyre^2^ engine^50^. The model was aligned to the *S. cerevisiae* sequence and visualized using the PyMOL Molecular Graphics System^51^.

### Ancestral character state reconstruction

Sequences of 240 fungal taxa were obtained from the Ensembl Genomes database for fungi and aligned with the murine homolog Hsp90ab1 (Table S4) in AliView version 1.26^52^ using the integrated default alignment program MUSCLE^53^. A Neighbor Joining tree was built in MEGA version 10.1.8^54^ and loaded into Mesquite version 3.61^55^ for ancestral character state reconstruction using the parsimony criterion.

### Western blotting and co-immunoprecipitations

To determine Hsp90 stability in the wild-type, *MAL2-HSP90* control and hsp90 mutant strains, stationary phase cells grown in YPM were inoculated into YPD or YPM at an OD_600_ of 0.2 and incubated until they reached a log-phase OD_600_ between 1.3 to 1.9 (approximately 4.5 hours). 50 ml of cell culture were pelleted, snap frozen in liquid nitrogen, and stored overnight at −20°C. Pellets were thawed on ice, washed twice with 1x PBS and resuspended in 250 μl lysis buffer (50 mM HEPES pH 7.5, 150 mM NaCl, 5 mM EDTA, 1% Triton X100, 1 cOmplete EDTA-free protease inhibitor tablet, 1 mM PMSF). Cells were transferred to Precellys 2 ml Tough Micro-organism Lysing Kit tubes (VK05) and lysed using a Precellys Evolution tissue homogenizer. Samples were processed in 3 cycles, each consisting of bead beating at 6,000 rpm for 1 minute, pause for 30 seconds, bead beating at 6,000 rpm for 1 minute. Tubes were rested on ice for at least 1 minute between each cycle. Lysates were cleared twice by centrifugation at 20,000 *x*g at 4°C for 30 minutes, and 5 minutes. Supernatants were transferred to microfuge tubes and refrigerated in preparation for sample preparation and gel loading. The lysates’ whole cell protein concentrations were determined with a Quick Start Bradford Protein Assay (Bio-Rad) and calibrated against a BSA standard curve. Samples were diluted to 0.1 μg/μl protein in sample buffer (70 mM Tris-HCl pH 6.8, 11.1% glycerol, 1.1% SDS, 0.005% bromophenol blue, 10% 2-mercaptoethanol, all final concentrations), heated at 95°C for 10 minutes and either cooled on ice prior to loading or stored at −20°C.

As loading controls, 100 μl per sample were loaded onto a 10% SDS-PAGE gel and separated for 20 minutes at 90 volts, followed by 100 minutes at 120 volts. To visualize proteins, gels were stained with SimplyBlue SafeStain (Invitrogen) according to the manufacturer’s instructions and imaged by scanning.

For immunoblotting of *C. albicans* Hsp90, 15 μl were loaded on a 10% SDS-PAGE gel and separated for 20 minutes at 90 volts, followed by 100 minutes at 120 volts. Proteins were wet-transferred onto a PVDF membrane (Bio-Rad) for 135 minutes at 300 mA in a Mini-PROTEAN Tetra Vertical Electrophoresis Cell filled with transfer-buffer (2 M glycine, 0.25 M Tris-base, 20% methanol). To block free sites, membranes were treated with PBST (1x PBS, 0.1% Tween 20) with 0.1% non-fat milk overnight at 4°C. The blots were probed for 1 hour at room temperature with α-CaHsp90^31^ diluted 1:10,000 in PBST with 0.1% milk and washed with 1x PBST prior to probing with α-rabbit antibody-HRP conjugate (BioRad #1705046) for 15 minutes at room temperature. Lastly, blots were washed with 200 ml PBST using the SNAP i.d. 2.0 Protein Detection System (Merck Millipore), incubated with Clarity Western ECL (BioRad), and imaged on the Syngene G:Box transilluminator.

To purify Hsp90 for mass-spectrometric analysis and to investigate the effect of Hsp90 phospho-mutant alleles on co-chaperone binding, 200 ml of cells were grown as described above, washed twice in 1x PBS and resuspended in 2 ml co-IP lysis buffer (50 mM Tris-HCl pH 7.5, 1% Nonidet P 40 Substitute (Sigma #74385), 0.25% deoxycholate Na, 150 mM NaCl, 5 mM EDTA, 1 mM Na_2_MoO_4_•2H_2_O, PhosSTOP tablet, cOmplete EDTA-free protease inhibitor tablet, 1 mM PMSF, 1 mM Na_3_VO_4_, and 10 mM NaF). Cells were transferred to 15 ml Precellys sample tubes (VWR BERT KT03961-1-4061), mixed with 1 ml acid-washed glass beads (~400 μm) and mechanically disrupted using a Precellys Tissue Homogeniser. Tubes were agitated three times for one minute with one minute on ice in between. The lysate was cleared by centrifugation at 20,000 *x*g twice for 5 minutes at 4°C and protein concentrations determined using the Bradford assay described above. 7.5 mg of protein were mixed with 100 μl of IgG or α-HA agarose, depending on the strain’s genotype, in co-IP washing buffer (50 mM Tris-HCl pH 7.5, 1% Nonidet P 40 substitute, 0.25% deoxycholate Na, 150 mM NaCl, 5 mM EDTA, 1 mM Na_2_MoO_4_•2H_2_O, 1 mM PMSF, 1 mM Na_3_VO_4_, and 10 mM NaF) and gently rotated for 2 hours at 4°C. Following precipitation, the agarose beads were washed carefully 3-5 times with 1 ml co-IP washing buffer before being mixed with 100 μl 1x sample buffer (6x sample buffer: 0.35 M Tris HCl, 10% (w/w) SDS, 36% glycerol, 5%2-mercaptoethanol and 0.012% bromophenol blue). Prior to loading, samples were heated at 95°C for 10 minutes and cooled on ice ready for loading or stored at −80°C. Proteins were separated at 120 volts on size-appropriate SDS-PAGE gels.

To visualize epitope-tagged co-chaperones, proteins were wet-transferred onto PVDF membranes for 16 hours at 20 volts in a Mini-PROTEAN Tetra Vertical Electrophoresis Cell filled with transfer-buffer (2 M glycine, 0.25 M Tris-base, 20% methanol). Membranes were blocked with PBST and 5% milk for at least 1 hour at room temperature before being probed with primary and secondary antibodies. To visualize TAP-tagged Sti1, membranes were treated with α-TAP antibody (Open Biosystems #CAB1001). Membranes carrying HA-tagged Cdc37, Aha1 or Sba1 were exposed to α-HA antibody (Invitrogen #71-5500). Both primary antibodies were diluted 1:5,000 in PBST with 0.2% milk and subsequently conjugated with α-rabbit antibody-HRP conjugate (BioRad Immun-Star Goat Anti-Rabbit (GAR)-HRP Conjugate #1705046) diluted 1:5,000. Membranes were incubated in antibody solutions for 15 minutes each and washed at least three times using 1x PBST using the SNAP i.d. 2.0 Protein Detection System. Lastly, proteins were visualized with Clarity Western ECL (BioRad).

### Temperature growth profiles

To assess susceptibility to thermal stress, optical densities of stationary phase cultures were measured at 600 nm and adjusted to 0.1 in 1x PBS and serially diluted two-fold. 4 μl of cell suspension were then spotted onto solid YPD or YPM and plates were incubated at the indicated temperature for 48 hours. Plates were photographed and analyzed visually.

### Cellular morphogenesis analysis

Stationary phase cells grown in YPM at 30°C were diluted to an optical density of 0.2 in YPD or YPM and grown for 16 hours at 30°C while shaking at 200 rpm. In preparation for microscopy, cells were washed and diluted with 1x PBS to a cell density appropriate for microscopy. Images were collected using the DIC settings on a Nikon Eclipse E8000 microscope with a Nikon Digital Sight DS-5M camera. To calculate the morphology index (MI)^*56*^, a total of 100 cells per strain were randomly selected and their width at the septal junction as well as cell length and maximum diameter measured using ImageJ^*57*^. The MI for each cell was then presented in scatter plots.

### Drug susceptibility assays

Strains were grown to stationary phase at 30°C in YPM while shaking at 200 rpm and their optical densities were measured at 600 nm and adjusted to ~10^3^ cells/100 μl. Fluconazole was diluted in water to a starting concentration of 512 μg/ml and 20μl of drug were mixed with 180 μl YPD or YPM in a 96-well flat bottom dish. The drug was then diluted two-fold in YPD or YPM to cover a gradient ranging from 256 μg/ml to 4 μg/ml. 100 μl of cell suspension were added to each well and plates were incubated for 48 hours at 25°C and optical densities recorded at 595 nm. A 5 mM radicicol stock solution (DMSO) was diluted to 50 μg/ml in YPM or YPD. The drug was diluted in either media to the following concentrations: 1, 2, 3, 4, 5, 7.5, 10, 15, 20, and 25 μg/ml and strains inoculated as above. To account for the effects of DMSO alone, strains were also inoculated in 0.0275% DMSO. Plates were incubated for 48 hours at 25°C in the dark and optical densities recorded at 595 nm. Optical density values were normalized to the no treatment column and visualized using Java TreeView version 1.1.6r4^*58*^.

### Colony morphology screen

Wild-type, control and mutant strains were grown overnight in YPM at 30°C while shaking at 200 rpm. Stationary cultures were then diluted to an optical density of 0.02 at 600 nm and 5 μl of cell suspension spotted onto RPMI, Spider, and synthetic defined media with and without Calcofluor White or Congo Red (Table S5). After incubation at 30°C for 48 hours colonies were visually scored for appearance as ‘smooth’, ‘rough’, ‘wrinkly’, or ‘wrinkly with hyphal edge’ and photographed.

### Biofilm analyses

In preparation for biofilm analyses, *C. albicans* strains were cultured overnight in YPD at 30°C while shaking at 150 rpm. Stationary phase cells were centrifuged at 4,400 rpm for 5 minutes, washed with 1x PBS and counted using a Neubauer hemocytometer. To induce biofilm formation, cells were diluted to 10^6^ cells/ml in RPMI (Sigma-Aldrich/Merck # R7755) and incubated at 37°C for 24 hours in flat-bottom 96-well dishes for cell viability and biomass assays or on Thermanox™ coverslips for scanning electron microscopy. Cell viability was assessed by metabolic reduction of resazurin sodium salt (Sigma-Aldrich/Merck #R7017). For the assay, 0.1 g of resazurin was dissolved in 10 ml 1x PBS and filter-sterilized. Biofilm wells were washed twice with 1x PBS and filled with 100 μl of a 1:1000 dilution of resazurin in RPMI. Plates were incubated at 37°C for 25 minutes and absorbance measured at 570 nm. Upon completion of the resazurin cell viability assay, plates were left to dry overnight and biofilm biomass assessed by staining with 0.05% w/v crystal violet. Upon washing and de-staining with 100% ethanol, biomass was quantified by spectrophotometric measurements at 570 nm in a FluoStar Omega (BMG Labtech) microtitre plate reader^*59*^. To visualize biofilms grown on coverslips, they were washed once with 1x PBS and fixed overnight at 4°C in 500 μl of 0.15% w/v Alcian Blue dissolved in 2% para-formaldehyde, 2% glutaraldehyde, and 0.15 M sodium cacodylate. Samples were imaged as described previously^*60*^.

### Virulence assays

To assess if Hsp90 phosphorylation affects virulence, we infected caterpillars of the Tobacco Hornworm *Manduca sexta*^38^, a novel invertebrate model of fungal disease. In preparation for virulence assays, *C. albicans* strains were grown overnight in YPD at 30°C and stationary phase cultures were washed twice with 1x PBS. Cell density was adjusted to 10^8^ cells/ml for each strain and groups of ten caterpillars were injected with 100 μl cell suspension per animal or 1x PBS in their left rear proleg. Caterpillars were maintained at 37°C and scored for survival after 16 hours.

### Statistical analyses

All analyses were carried out in R Version 3.6.0 and plotted using ggplot2. To assess the effect of the Hsp90 mutant alleles on cellular morphology, the MIs for samples grown in YPD and YPM were compared with each other using the Mann-Whitney U test. Statistical differences between biofilm cell viability and biomass production were assessed using a linear model of strain versus optical density comparing the mutant strains to the maltose control. Statistical significance of caterpillar survival was assessed using Fisher’s exact test.

## Acknowledgments

We would like to thank Prof Alistair J. P. Brown for comments on the manuscript, Ewan Basterfield and Chris Apark for technical support with the *Manduca* caterpillars, and Profs Julia R. Köhler, Jürgen Wendland, and Malcolm Whiteway for their generous plasmid gifts. This work was funded by MRC grant MR/L018349/1 to SD. JLC was supported by the MRC GW4 ‘Biomed’ Doctoral Training Program.

## Author contributions

LA, JLC and SD conceived and designed this study. LA constructed strains, prepared samples for mass spectrometry, conducted Hsp90 stability assays and assessed co-chaperone associations. JLC conducted Hsp90 stability and drug susceptibility assays. UO performed the structural modelling analysis. RW performed Hsp90 phylogenetic analyses and ancestral character state reconstruction. CEW performed *in vivo* assays. HOB analyzed growth, morphology, and biofilm data. CS assisted with mass spectrometric analyses. MB & GR performed and analyzed biofilm formation. SD drafted and edited the manuscript.

## Additional information

Supplementary information is available for this paper.

Correspondence and requests for materials should be addressed to SD.

Code availability: All custom code is available from: https://github.com/hobrien/DiezmannLab/blob/master/T25_analysis.Rmd

