## Supplementary material for "A novel Hsp90 phospho-switch modulates virulence in the major human fungal pathogen *Candida albicans*": Alaalm Supplementary Materials (from another version of this preprint: version 1, 2020-09-23)

### SUPPLEMENTARY INFORMATION

#### Supplementary Tables

**Table S1: Strains used in this study, their progenitor strains and the plasmids necessary in strain construction.**

| Strain Name | Strain ID | Genotype | Progenitor | Plasmid |
|---|---|---|---|---|
| SN95[1] | YSD89 | *arg4Δ/arg4Δ his1Δ/his1Δ URA3/ura3Δ::imm$^{434}$ IRO1/iro1Δ:: imm$^{434}$* | | |
| Mal2-Hsp90 | YSD1384 | As SN95 *HIS1-MAL2p-HSP90/HSP90* | YSD89 | pSD94* |
| CK2 subunit deletion strains with TAP-tagged Hsp90 | | | | |
| cka1 | YSD557 | As SN95 *cka1Δ/Δ* | YSD89 | pSD3 |
| cka1_Hsp90-TAP | YSD692 | As SN95 *cka1Δ/Δ HSP90/HSP90-TAP-HIS1* | YSD557 | pSD18 |
| cka2 | YSD623 | As SN95 *cka2Δ/Δ* | YSD89 | pSD3 |
| cka2_Hsp90-TAP | YSD675 | As SN95 *cka2Δ/Δ HSP90/HSP90-TAP-HIS1* | YSD623 | pSD18 |
| ckb1 | YSD628 | As SN95 *ckb1Δ/Δ* | YSD89 | pSD3 |
| ckb1_Hsp90-TAP | YSD694 | As SN95 *ckb1Δ/Δ HSP90/HSP90-TAP-HIS1* | YSD628 | pSD18 |

| Strain Name | Strain ID | Genotype | Progenitor | Plasmid |
|---|---|---|---|---|
| ckb2 | YSD634 | As SN95 *ckb2Δ/Δ* | YSD89 | pSD3 |
| ckb2_Hsp90-TAP | YSD696 | As SN95 *ckb2Δ/Δ HSP90/HSP90-TAP-HIS1* | YSD634 | pSD18 |
| Non-phosphorylatable T25A allele construction | | | | |
| T25A-TAP | YSD1083 | As SN95 *HSP90/hsp90*$^{T25A}$*-TAP-ARG4* | YSD89 | pSD72 |
| Mal2-Hsp90 T25A-TAP | YSD1386 | *HIS1-MAL2p-HSP90/hsp90*$^{T25A}$*-TAP-ARG4* | YSD1083 | pSD94 |
| Mal2-Hsp90 T25A-NAT | YSD1398 | *HIS1-MAL2p-HSP90/hsp90*$^{T25A}$*-NAT-FLP* | YSD1386 | pSD3 |
| Mal2-Hsp90 T25A | YSD1403 | *HIS1-MAL2p-HSP90/hsp90*$^{T25A}$ | YSD1398 | |
| Phosphomimetic T25E allele construction | | | | |
| T25E-TAP | YSD1085 | As SN95 *HSP90/hsp90*$^{T25E}$*-TAP-ARG4* | YSD89 | pSD74 |
| Mal-Hsp90 T25E-TAP | YSD1389 | *HIS1-MAL2p-HSP90/hsp90*$^{T25E}$*-TAP-ARG4* | YSD1085 | pSD94 |
| Mal-Hsp90 T25E-NAT | YSD1402 | *HIS1-MAL2p-HSP90/hsp90*$^{T25E}$*-NAT-FLP* | YSD1389 | pSD3 |
| Mal-Hsp90 T25E | YSD1410 | *HIS1-MAL2p-HSP90/hsp90*$^{T25E}$ | YSD1402 | |

| Strain Name | Strain ID | Genotype | Progenitor | Plasmid |
|---|---|---|---|---|
| Non-phosphorylatable S530A allele construction | | | | |
| S530A-TAP | YSD1434 | As SN95 *HSP90/hsp90*$^{S530A}$*-TAP-ARG4* | YSD89 | pSD97 |
| Mal2-Hsp90 S530A-TAP | YSD1437 | *HIS1-MAL2p-HSP90/hsp90*$^{S530A}$*-TAP-ARG4* | YSD1434 | pSD94 |
| Mal2-Hsp90 S530A-NAT | YSD1440 | *HIS1-MAL2p-HSP90/hsp90*$^{S530A}$*-NAT-FLP* | YSD1437 | pSD3 |
| Mal2-Hsp90 S530A | YSD1442 | *HIS1-MAL2p-HSP90/hsp90*$^{S530A}$ | YSD1440 | |
| Phosphomimetic S530E allele construction | | | | |
| S530E-TAP | YSD1435 | As SN95 *HSP90/hsp90*$^{S530E}$*-TAP-ARG4* | YSD89 | pSD98 |
| Mal-Hsp90 S530E-TAP | YSD1439 | *HIS1-MAL2p-HSP90/hsp90*$^{S530E}$*-TAP-ARG4* | YSD1435 | pSD94 |
| Mal-Hsp90 S530E-NAT | YSD1441 | *HIS1-MAL2p-HSP90/hsp90*$^{S530E}$*-NAT-FLP* | YSD1439 | pSD3 |
| Mal-Hsp90 S530E | YSD1444 | *HIS1-MAL2p-HSP90/hsp90*$^{S530E}$ | YSD1441 | |
| Epitope-tagged co-chaperones in SN95 and phospho-mutant strains | | | | |
| Sti1-TAP | YSD440 | As SN95 *STI1/STI1-TAP-ARG4* | YSD89 | pSD19 |

| Strain Name | Strain ID | Genotype | Progenitor | Plasmid |
|---|---|---|---|---|
| Aha1-HA | YSD1244 | As SN95 *AHA1/AHA1-HA-ARG4* | YSD89 | pSD22 |
| Sba1-HA | YSD1246 | As SN95 *SBA1/SBA1-HA-ARG4* | YSD89 | pSD22 |
| Cdc37-HA | YSD826 | As SN95 *CDC37/CDC37-HA-ARG4* | YSD89 | pSD22 |
| Sti-TAP Mal-Hsp90 T25A | YSD1423 | *STI1/STI1-TAP-ARG4 HIS1-MAL2p-HSP90/hsp90*$^{T25A}$ | YSD1403 | pSD19 |
| Aha1-HA Mal-Hsp90 T25A | YSD1419 | *AHA1/AHA1-HA-ARG4 HIS1-MAL2p-HSP90/hsp90*$^{T25A}$ | YSD1403 | pSD22 |
| Sba1-HA Mal-Hsp90 T25A | YSD1421 | *SBA1/SBA1-HA-ARG4 HIS1-MAL2p-HSP90/hsp90*$^{T25A}$ | YSD1403 | pSD22 |
| Cdc37-HA Mal-Hsp90 T25A | YSD1422 | *CDC37/CDC37-HA-ARG4 HIS1-MAL2p-HSP90/hsp90*$^{T25A}$ | YSD1403 | pSD22 |
| Sti-TAP Mal-Hsp90 T25E | YSD1431 | *STI1/STI1-TAP-ARG4 HIS1-MAL2p-HSP90/hsp90*$^{T25E}$ | YSD1410 | pSD19 |
| Aha1-HA Mal-Hsp90 T25E | YSD1425 | *AHA1/AHA1-HA-ARG4 HIS1-MAL2p-HSP90/hsp90*$^{T25E}$ | YSD1410 | pSD22 |
| Sba1-HA Mal-Hsp90 T25E | YSD1428 | *SBA1/SBA1-HA-ARG4 HIS1-MAL2p-HSP90/hsp90*$^{T25E}$ | YSD1410 | pSD22 |
| Cdc37-HA Mal-Hsp90 T25E | YSD1429 | *CDC37/CDC37-HA-ARG4 HIS1-MAL2p-HSP90/hsp90*$^{T25E}$ | YSD1410 | pSD22 |

*previously published plasmids:
pSD3 (NAT1-flipper pJK863[2])

| Strain Name | Strain ID | Genotype | Progenitor | Plasmid |
|---|---|---|---|---|

pSD18 (pFA-TAP-HIS1)[3]
pSD19 (pFA-TAP-ARG4)[3]
pSD22 (pFA-HA-ARG4)[3]
pSD94 (pFA-CaHIS1-MAL2p)[4]

plasmids pSD72, pSD74, pSD97, and pSD98 – this study.

**Table S2:** Primers for strain and plasmid construction.

| Primer Name | Primer ID | Primer Sequence (5' to 3') |
|---|---|---|
| Insertion of *MAL2*p-ON promoter upstream of Hsp90 | | |
| CaHSP90_*MAL2*p_-118F | oSD845 | TTGCCAATATATAAATTTACGTGAAAAATTCTTCACCAGTTTATCAACT CCTCCCGTTTTTCTTTTTTCATTTCTTCTTTCTATCCAAATCATACC**G AAGCTTCGTACGCTGCAGGTC** |
| CaHSP90_*MAL2*p_+92R | oSD883 | TCCTTGTTTGAATAGACTGTGTTAATGATCAAAGACATCAACTGAGAG ATCTCAGCAGTGAATTCGTGAGTTTCAACTTTTGCGTCAGCCAT**CATT GTAGTTGATTATTAGTTAAACCAC** |
| Construction of CK2 gene deletion mutants and Hsp90 TAP-tagging | | |
| CKA1-U100F | oSD424 | GTTGTTGTATCCATTTTTAATACCAGTATCCTTTTTGACTTTGCTAATTTCCTAT CGCTATTCATTGCCACATAACAAACTTTTTAATAGCAATAGCACA**GGAAACA GCTATGACCATG** |
| CKA1-D100R | oSD425 | AGTATGTAGAACCACACGCCATCATCGTAACACATTTTTTTCATGAAATTTAC ATTTCTATATAAAAATATTACAGCAATACAAAAAATAACGGGCAGACTTA**GT AAAACGACGGCCAG** |
| CKA2-U100F | oSD428 | CAGATTGGATTAATATATTTAAATAAACTTTATTCCTCAATTAATCCATATTAT ATATCCATATATATTTATACACACATATCAATAATTATTATTAGTA**GGAAACA GCTATGACCATG** |
| CKA2-D100R | oSD429 | TAGTAAGAGAAAATAAATAAATAGTTTTAATGTCGGTTGTTAATTGTGTTTGA GACATTTTATGCAGCAATGACTAATTGTATATATAATAAATTATATACTT**GTA AAACGACGGCCAG** |
| CKB1-U100F | oSD416 | TGTCACCACTAGTTTTTTTTTTTTTACCTCCATTGATAAAACAAGAACGGAAA AAAAAAAAATTAAGATTAACATAGCTTTGCCGTGATATCAACTACA**GGAAAC AGCTATGACCATG** |

| Primer Name | Primer ID | Primer Sequence (5’ to 3’) |
|---|---|---|
| CKB1-D100R | oSD417 | TCATTAAATAAGCTATTATTTACCTCTTTATATTTATTATTATACACGAAAAGA TGTACACACACACATATACGTCAATTATTCTTTCCTTTTATACGTTTCT**GTAAA ACGACGGCCAG** |
| CKB2-U100F | oSD420 | GTTTTTATTTTTAAACTTCAGTTAATATTAAATATACCAAATAACTAATTCTTC TATTTTTTTTTTGTTTTATTTCAGATTTACTAAGTTATAACAAAAG**GGAAACAG CTATGACCATG** |
| CKB2-D100R | oSD421 | TTTTCTTTTTCATTTATTTATTAATACCAATATAGCAGCAAAAAAGTTTATTGA AGTACTCTAGTATTTACAAAACTGGTTGTGGTAGTGGTGGGACGTGATG**GTAA AACGACGGCCAG** |
| Ca-HSP90-TAP-F | oSD84 | TGAACCTGAAGCTACTACTACTGCCTCAACTGACGAACCAGCTGGAGAATCTG CTATGGAAGAAGTTGAT**GGTCGACGGATCCCCGGGTT** |
| Ca-HSP90-TAP-R | oSD85 | ATGTTATTACTCTCTAGATACACGATATTACAAAACTTATTTAACTAGAAAAC TGTAGCCCTTCTGGTGT**TCGATGAATTCGAGCTCGTT** |
| Construction of mutagenized Hsp90 alleles | | |
| CaHSP90_-120F_TAP | oSD712 | GGTTGCCAATATATAAATTTACGTGAAAAATTCTTCACCAGTTTATCA ACTCCTCCCGTTTTTTCTTTTTTTCATTTCTTCTTTCTATCCAAATCATA CCATACAAATCAATAGTTCATT**GGTCGACGGATCCCCGGGTT** |
| CaHSP90_+2241R_TAP | oSD713 | TCGATTTAATGTACTTTATCTCTTTCTAGAAAAAAGAACCATAATTCA ATAATACAAGCCAAGTCTCGAGATTTTTCGATTTTCCTTTTAGCATGCG CAT**TAGTGGATCTGATATCATCG** |
| CaHSP90_+2023F | oSD773 | TTGACGATGATTCAGAAGAAACTGCTGTTGAACCTGAAGCTACTACTA CTGCCTCAACTGACGAACCAGCTGGAGAATCTGCTATGGAAGAAGTT GATTAA**GGAAACAGCTATGACCATG** |
| CaHSP90_+2344R | oSD774 | TCGATTTAATGTACTTTATCTCTTTCTAGAAAAAAGAACCATAATTCA ATAATACAAGCCAAGTCTCGAGATTTTTCGATTTTCCTTTTAGCATGCG CATTGAG**GTAAAACGACGGCCAG** |

| Primer Name | Primer ID | Primer Sequence (5' to 3') |
| --- | --- | --- |
| CaHSP90_MET3p_-118F | oSD845 | TTGCCAATATATAAATTTACGTGAAAAATTCTTCACCAGTTTATCAACTCCTCCCGTTTTTCTTTTTTTCATTTCTTCTTTCTATCCAAATCATACC**GAAGCTTCGTACGCTGCAGGTC** |
| CaHSP90_MAL2p_+92R | oSD883 | TCCTTGTTTGAATAGACTGTGTTAATGATCAAAGACATCAACTGAGAGATCTCAGCAGTGAATTCGTGAGTTTCAACTTTTGCGTCAGCCAT**CATTGTAGTTGATTATTAGTTAAACCAC** |
| Co-chaperone TAP-tagging | | |
| STI1_+1698_TAP_F | oSD229 | TCATATGAAAAATCCTGAAGTTTATAAAAAAATTAATATGTTGATTGCTGCTGGTGTTATTCGTACCAGA**GGTCGACGGATCCCCGGGTT*** |
| STI1_+1854_TAP_R2 | oSD242 | CAGTTTGATATATATAATATATATATTTGCCTATGTATATGCTAACTAATTATATGACACCAAAAATAGA**TCGATGAATTCGAGCTCGTT** |
| AHA1-HA-F | oSD769 | AGTGGAATCCCAATTGGTGAAGAGGAAAGAGTGAAAAATAACTTTGAAGAAAGATATATCAGATCTATAAAGATCACTTTTGGATTTGGAGCAGTTTTA**TACCCATACGATGTTCCTGAC** |
| AHA1_TAP_R | oSD210 | GGTATCATAAGTGATGGTTAATAGAGTAAAATTCTTCTTTATGGATTGCTTGTACGTACATTCTATTTAT**TCGATGAATTCGAGCTCGTT** |
| SBA1_HA_F | oSD768 | GGTGGTGCTGCAGGTCTTGATGGCGAAGAAGGCGAAGAAGGCGAAGCAGAAGCTAAAGAAGCGCAAGAAGAATCAAATACCACTGCTACTGAAAAAGAA**TACCCATACGATGTTCCTGAC** |
| SBA1_TAP_R | oSD226 | AAATGACTAATAAAATAGACTATTCAACTGTATAATTTTGGTTTTTTTTGTATTACTTTGTTATTTGAGA**TCGATGAATTCGAGCTCGTT** |
| CDC37-HA-F | oSD656 | GAAGATGAAAAGGAATTTGAAGAATTGAAAAAAGAATATGCACATGAAACTGCCAATCAAGAAGAAGACCAGTCTGCTTCAGTTGAAGATACAGTTGAT**TACCCATACGATGTTCCTGAC** |

| Primer Name | Primer ID | Primer Sequence (5’ to 3’) |
| --- | --- | --- |
| Ca-CDC37-TAP-R | oSD61 | ATATCCCCTGAAAAAAAACCACGAAATCTACAATTCGACTAGGGTAC AAGTGCAATTTTTATTACGCTAT**TCGATGAATTCGAGCTCGTT** |
| T25 mutagenic plasmids (pSD72 and pSD74) | | |
| CaHSP90_NotI_-128F | oSD597 | CGAGCGGCCGCTCTCCGTTGGTTGCCAATAT |
| CaHSP90_Not1_+2293R | oSD598 | CGAGCGGCCGCATACAAGCCAAGTCTCGAGA |
| CaHSP90_T25A-F | oSD642 | GTCTTTGATCATTAACGCAGTCTATTCAAACAAGG |
| CaHSP90_T25A-R | oSD643 | CCTTGTTTGAATAGACTGCGTTAATGATCAAAGAC |
| CaHSP90_T25E-F | oSD644 | GTCTTTGATCATTAACGAAGTCTATTCAAACAAGG |
| CaHSP90_T25E-R | oSD645 | CCTTGTTTGAATAGACTTCGTTAATGATCAAAGAC |
| CaHSP90_-133F_PacI | oSD708 | TTAATTAAAGTGGTCTCCGTTGGTTGCCA |
| CaHSP90_+2100R_NcoI | oSD709 | CCATGGATCAACTTCTTCCATAGCAGA |

[2*] bold sequences indicate homology to plasmid DNA.

**Table S3:** Diagnostic PCR and sequencing primers used for strain and plasmid construction.

| Primer Name | Primer ID | Primer Sequence (5’ to 3’) |
|---|---|---|
| TAP_R | oSD33 | TAAACTTTGGATGAAGGCG |
| ARG4-F | oSD34 | ATGTTGGCTACTGATTTAGCTG |
| Ca-CDC37-+983F | oSD62 | AAGCAGCACCAGCTAATGTG |
| Ca-HSP90-+1617F | oSD86 | GAAATCAAAGAATACGAACCAT |
| STI1_+1474F | oSD231 | AGGCAAGAACTAAAGATGTTG |
| STI1_+1877R | oSD232 | TTTGCCTATGTATATGCTAAC |
| pJK863down-F | oSD317 | CTGTCAAGGAGGGTATTCTGG |
| pJK863up-R | oSD318 | AAAGTCAAAGTTCCAAGGGG |
| Ca_HSP90_+511R | oSD383 | CTCAACATGGTACCACGACC |
| CKB1-UpF | oSD418 | GGTAGAACTCGTCTCTGGTG |
| CKB2-UpF | oSD422 | GGGTAAATTGGATGGGTCGC |
| CKA1-UpF | oSD426 | CAGCAAGCTGCCAGCTCAGT |
| CKA2-UpF | oSD430 | GTGATAGTAAATCTCCTTGG |
| CKA1-DownR | oSD450 | GGAGTGTGGACGAGTAGTGT |
| CKA2-DownR | oSD451 | GAAGCTCGTGATCCATCTGA |
| CKB2-DownR | oSD453 | CCAGCAGCTGTGGTTAATGG |
| CKB1-DownR-2 | oSD463 | TCCATATACTCGAATTGCAC |
| TAP-R3 | oSD522 | AGGCGTTTCGTTGTTCTTCG |
| Ca_HSP90_+308F | oSD529 | GTACCAAATCCTTTATGGAAGC |
| Ca_HSP90_+1002F | oSD530 | GAAGAACAACATCAAATTATAC |
| CaHSP90_+1042F | oSD554 | CTGATGATGCTGAAGAGTTG |
| CaHSP90_+1132R | oSD555 | CTCTGGACAAGTTCAATGGC |

| | | |
|---|---|---|
| pSD3-NotI-UpF | oSD601 | GTAATACGACTCACTATAGG |
| pSD3-Not1-DownR | oSD602 | CTAGAGAATAGGAACTTCAG |
| CaHSP90_+612F | oSD611 | GCTTATCCAATCCAATTAGT |
| CaHSP90-+1292R | oSD612 | TGAGCATCTTCATGAATACC |
| HA-R | oSD652 | ATGGATATCCTGCATAGTCC |
| CaHSP90_+2710R | oSD654 | GTGACAAGCTATGACATGAC |
| CaHSP90_-169F | oSD723 | GGTTCTATAGAATTCCATCA |
| SBA1_+46F | oSD794 | GTTCATCTGAAGATGACGCT |
| AHA1_+451F | oSD795 | CACAAGATAAGGTCACCTCG |
| Ca_STI1_+1228F | oSD796 | TCACCAAAGGAGATTGGCCA |
| MAL2p-F | oSD880 | GTTTAAACTTTTGTCTAGTACCATCTGTACC |
| CaHSP90_+1051F | oSD886 | GCTGAAGAGTTGATTCCAGAAT |

**Table S5:** Media for morphology screen.

| Media | Component |
|---|---|
| RPMI | 1.04% RPMI 1640<br>3.425% MOPS<br>2% dextrose<br>0.008% uridine<br>0.002% histidine<br>0.01% leucine<br>1.5% agar |
| Spider | 1% nutrient broth<br>1% mannitol<br>0.2% dipotassium phosphate<br>0.008% uridine<br>1.35% agar |
| Synthetic defined (SD) | 0.17% yeast nitrogen base<br>0.5% ammonium sulphate<br>2% dextrose<br>1.5% agar |
| Congo Red | SD + 0.02% Congo Red |
| Calcofluor White | SD + 0.015% Calcofluor White M2R |

**Table S6:** Survival of *M. sexta* caterpillars 16 hours post infection with $10^7$ *Candida* cells.

| Treatment | # survivors | p-value |
|---|---|---|
| PBS | 10 | 0.0007145 |
| WT | 4 | 0.628483 |
| *MAL2-HSP90* | 2 | |
| T25A | 5 | 0.3498452 |
| T25E | 7 | 0.0697785 |
| S530A | 7 | 0.0697785 |
| S530E | 9 | 0.0054775 |

### Supplementary Methods

#### Plasmid construction

**pSD72 (T25A) mutagenic plasmid.** To generate pSD51 (pSD3-CaHSP90-NAT), the *HSP90* ORF was PCR amplified from YSD91 (SN78) gDNA using oligonucleotides oSD597 and oSD598. The amplicon was digested with NotI and ligated into NotI linearized pSD3 (pJK863; SAP2-FLP2-NAT1[2]). Successful integration was verified by genotyping PCR of the HSP90 ORF with oligonucleotides oSD601 and oSD602 and by validation of correct upstream (oSD601 and oSD383) and downstream (oSD602 and oSD86) integration. The construct was furthermore validated with a diagnostic NotI digest and Sanger sequencing using primers oSD554, oSD555, oSD601, oSD602, oSD611, and oSD612.

To generate pSD59 (pSD3-hsp90-T25A-NAT), pSD51 was used as template to amplify the cassette with the mutagenic oligonucleotides oSD642 and oSD643 thereby introducing the T25A mutation into *HSP90*. After treating the PCR product with Dpn1 to digest the wild-type template, the mutagenized plasmid was transformed into *Escherichia coli* DH5α cells and sequence verified with oSD383.

To generate pSD68 (pFA-CaHSP90-TAP-ARG4), the *HSP90* ORF excluding the stop codon was PCR amplified from SN78 (YSD91) gDNA using the primer pair oSD708 and oSD709, which also integrated Pac1 and NcoI restriction sites. The double digested insert was then ligated into linearized double digested pSD19 (pFA-TAP-ARG4[3]) and successful integration was verified by diagnostic PacI and Ncol digests before being sequenced using primers oSD383, oSD529, oSD530, and oSD86.

To generate pSD72 (pFA-hsp90-T25A-TAP-ARG4), pSD59 was used to amplify the non-phosphorylatable *hsp90*$^{T25A}$ allele using primers oSD708 and oSD709, which also served to integrate PacI and NcoI restriction sites. The product was sequence verified by using oSD383 and cloned into pSD19 (pFA-TAP-ARG4[3]). The mutagenized plasmid was transformed into *E. coli* cells and verified by sequencing using primers oSD86, oSD383, oSD529, and oSD530.

**pSD74 (T25E) mutagenic plasmid.** To generate pSD74 (pFA-hsp90-T25E-TAP-ARG4), pSD68 (pFA-HSP90-TAP-ARG4), carrying wild type *HSP90*, was used as a template to generate the mutant *hsp90*$^{T25E}$ allele. To do so, *HSP90* was amplified off pSD68 with two sets of primers, oSD708 and oSD644 as well as oSD645 and oSD709. This introduced the required point mutation and integrated PacI and Ncol restriction sites. The overlapping PCR products were then amplified with oSD708 and oSD709 to yield full-length *hsp90*$^{T25E}$. This product was verified by sequencing using oSD383, before being cloned into pSD19 (pFA-TAP-ARG4[3]). The mutagenized plasmid was transformed into *E. coli* and verified by sequencing using oSD86, oSD383, oSD529 and oSD530.

**pSD97 and pSD98 mutagenic plasmids.** Both plasmids were generated by GenScript based on the backbone of plasmid pSD68 (pFA-*HSP90*-TAP-ARG4), which was provided. Following gene synthesis, pSD97 carries the non-phosphorylatable S530A allele and pSD98 the phosphomimetic S530E allele.

***Candida albicans* strain construction**

To construct strains carrying the *MAL2* inducible promoter, *HSP90* phospho-mutant alleles, or epitope tags, SN95 and its derivatives were transformed with PCR amplicons off the appropriate plasmids (Table S1). Amplicons were generated by PCR amplification of 0.1 ng/µl of the appropriate plasmid with primers with 100 bp homology to the genomic target locus (Table S2). 0.4 µM forward and reverse primer were mixed with 2 U of Q5 High Fidelity DNA polymerase (NEB), 5x reaction buffer, and 25 µM dNTPs in a total volume of 50 µl. Amplification started with an initial denaturation at 98˚C for 2 minutes, followed by 30 cycles of denaturation at 98˚C for 30 seconds, annealing at 55˚C for 30 seconds, elongation at 72˚C for 3 minutes 30 seconds, and concluded with a final elongation step at 72˚C for 2 minutes. Successful amplification was verified by electrophoretic separation on 1% agarose gels. Lastly, 400 µl PCR product were precipitated with 95% ethanol, pellets washed once with 70% ethanol and dissolved in 50 µl distilled $H_2O$ prior to DNA transformation.

*C. albicans* cells were transformed using the lithium-acetate protocol[5]. Briefly, each strain was grown overnight in 10 ml YPD at 30˚C while shaking at 200 rpm to a target OD between 4 and 8. To calculate how much of the cell suspension was needed for each transformation reaction, the OD was multiplied by 1.5, yielding the number of possible transformations from a 10 ml cell culture volume. The required amount of cells was washed once with sterile water and transformation mix (800 µl 50% sterile-filtered PEG 3350, 100 µl 10x TE buffer, 100 µl 1 M lithium acetate pH 7.4, 20 µl 1 M 1,4 dithiothreitol, 40 µl 5 mg/ml denatured salmon sperm DNA, ethanol purified PCR product) was added. Cells were gently mixed and incubated at 30˚C for 1 hour before being carefully transferred to 42˚C for 20 minutes.

Following incubation, transformation reactions were centrifuged for 15 secs and the transformation mix was removed. Cells transformed with the arginine or histidine auxotrophic marker where resuspended in 600 µl sterile distilled water and gently spread onto selective plates (DIFCO, 0.67% yeast nitrogen base without arginine or histidine, 2% glucose). Cells transformed with the NAT marker were resuspended in 600 µl YPM, incubated at 30˚C while shaking for at least 3 hours, and spread onto YPM plates containing 150 µg/ml nourseothricin. Plates were incubated for 3 to 5 days at 30˚C. All colonies were re-streaked onto selective media before being genotyped for correct allele integration by PCR amplification across the up- and downstream integration sites.

For diagnostic PCR, a minute amount of yeast cells was mixed with 2x PCR master mix (Quantig Ltd.) and 0.2 µM of forward and reverse primer (Table S3) in a total volume of 25 µl. PCR reactions were incubated for an initial denaturation at 94°C for 5 minutes, followed by 30 cycles of denaturation at 94°C for 30 seconds, annealing at 53°C for 45 seconds and elongation at 72°C for 1 minute. The program concluded with a final elongation at 72°C for 10 minutes. PCR products were then separated on a 1% agarose gel and screened for amplicons indicative of correct integration. Following confirmation of correct allele integration, the NAT marker was excised by incubation of the $NAT^R$ positive transformant in 5 ml YNB-BSA (0.85 g YNB without amino acids and ammonium sulfate, 2 g bovine serum albumin, 1 g yeast extract, 10 g maltose in 500 ml $H_2O$) for at least two days at 30˚C. The cell culture was diluted and spread onto appropriate media without nourseothricin from where colonies were replica plated onto media with 150 µg/ml nourseothricin for identification of $NAT^S$ colonies.

**T25 mutant strain construction and epitope tagging.** To generate the TAP-tagged T25 heterozygous mutants, YSD1083 (HSP90/hsp90-T25A-TAP-ARG4) and YSD1085 (HSP90/hsp90-T25E-TAP-ARG4), the cassettes were PCR amplified from pSD72 and pSD74 using primers oSD712 and oSD713. Amplicons were transformed into YSD89 (SN95) and transformants genotyped using primers oSD33 and oSD723. The mutant allele was further validated by sequencing using primers oSD86, oSD383, oSD529, oSD530, oSD611, and oSD612 primers after having been amplified with oSD33 and oSD723 from genomic DNA using Q5 DNA polymerase.

To control the expression of wild type *HSP90* with *MAL2*p in the T25 heterozygous mutants and to generate YSD1386 (HIS1-MAL2p-HSP90/hsp90-T25A-TAP-ARG4) and YSD1389 (HIS1-MAL2p-HSP90/hsp90-T25E-TAP-ARG4), the *HIS1*-*MAL2*p cassette was amplified with primers oSD845 and oSD883 from pSD94. Amplicons were transformed into T25 heterozygous mutants YSD1083 (HSP90/hsp90-T25A-TAP-ARG4) and YSD1085 (HSP90/hsp90-T25E-TAP-ARG4). Transformants were genotyped with primer pair oSD880 and oSD383 amplifying *MAL2*p and primer pair oSD86 and oSD522 amplifying the TAP tag. To ensure that *MAL2*p and TAP are not in the same allele, transformants were screened with primers oSD880 and oSD522.

To replace the TAP-ARG4 cassette with the recyclable NAT marker, the NAT cassette was PCR amplified with primers oSD773 and oSD774 from linearized plasmid pSD3. The amplicon was transformed into strains YSD1386 (HIS1-MAL2p-HSP90/hsp90-T25A-TAP-ARG4) and YSD1389 (HIS1-MAL2p-HSP90/hsp90-T25E-TAP-ARG4) and transformants were genotyped for correct upstream integration with primers oSD86 and oSD318 and correct downstream integration with primers oSD654 and oSD317. Before flipping out the NAT marker, transformants were tested for arginine requirements to make sure that NAT had not inserted into *MAL2*p-*HSP90*. Positive transformants only grew on YNB with arginine but not on YNB without arginine.

To generate YSD1403 (HIS1-MAL2p-HSP90/hsp90-T25A) and YSD1410 (HIS1-MAL2p-HSP90/hsp90-T25E), the NAT marker was flipped out from YSD1398 (HIS1-MAL2p-HSP90/hsp90-T25A-NAT) and YSD1402 (HIS1-MAL2p-HSP90/hsp90-T25E-NAT) as described above.

**S530 mutant construction.** To generate strains carrying the phosphomimetic and non-phosphorylatable *HSP90*$^{S530}$ alleles, the HSP90*-TAP-ARG4 cassettes were amplified off plasmids pSD97 and pSD98 using primers oSD712 and oSD713 and transformed into SN95 to generate strains YSD1434 (S530A-TAP) and YSD1435 (S530E-TAP). Positive integration and replacement of one HSP90 wild-type allele was verified by genotyping PCR using primers oSD33 and oSD723. The construct was then amplified from *C. albicans* gDNA using oSD33 and oSD723 and sequence validated using primers oSD86, oSD383, oSD529, oSD611 and oSD886.

Following successful replacement of one *HSP90* wild-type allele with the mutant allele, the remaining wild-type allele was placed under the control of the *MAL2*p promoter by amplifying the *HIS1-MAL2*p cassette from pSD94 using primers oSD845 and oSD883 and transformation into the *HSP90*$^{S530}$ heterozygous mutants YSD1434 (*HSP90/hsp90*$^{S530A}$*-TAP-ARG4*) and YSD1435 (*HSP90/hsp90*$^{S530E}$*-TAP-ARG4*). Transformants were genotyped with primers oSD880 and oSD383 amplifying *MAL2*p and primers oSD86 and oSD522 amplifying the TAP tag. To ensure that *MAL2*p and TAP are not in the same allele, transformants were screened with primers oSD880 and oSD522.

Strains YSD1437 (HIS1-MAL2p-HSP90/hsp90$^{S530A}$-TAP-ARG4) and YSD1439 (HIS1-MAL2p-HSP90/hsp90$^{S530E}$-TAP-ARG4) were then transformed with the NAT cassette replacing the *TAP-ARG4* construct. The NAT cassette was amplified using primers oSD773 and oSD774 from linearized plasmid pSD3 (pJK863[2]) and transformants screened for successful integration with primer pairs oSD86 and oSD318 as well as oSD654 and oSD317. This yielded strains YSD1440 (Mal2-Hsp90 S530A-NAT) and YSD1441 (Mal-Hsp90 S530E-NAT). Before flipping out the NAT marker, transformants were tested for arginine requirements to make sure that NAT had not inserted into *MAL2*p-*HSP90*. Positive transformants only grew on YNB with arginine but not on YNB without arginine. Removing the NAT marker then yielded strains YSD1442 (Mal2-Hsp90 S530A) and YSD1444 (Mal-Hsp90 S530E).

**Epitope tagging of Cdc37, Sba1, Aha1 and Sti1 in T25 mutants.** Select co-chaperones were C-terminally epitope-tagged in the arginine auxotroph T25 mutants YSD1403 (HIS1-MAL2p-HSP90/hsp90-T25A) and YSD1410 (HIS1-MAL2p-HSP90/hsp90-T25E). Cdc37, Sba1, Aha1 were HA-tagged and Sti1 was TAP-tagged. To do so, the TAP-ARG4 and HA-ARG4 cassettes were PCR amplified from 0.1 ng/µl of pSD19 (pFA-TAP-ARG4) and pSD22 (pFA-HA-ARG4) with primers with 100bp homology to the target genes (Table S2) using Q5 Taq Polymerase. Arginine prototroph transformants were genotyped using primers specific to the co-chaperone (AHA1: oSD795, Sba1: oSD794, Cdc37: oSD62, Sti1: oSD796) and the epitope (TAP: oSD522, HA: oSD652) (Table S2).

To construct Hsp90-epitope tagged CK2 deletion mutants for mass spectrometric analysis of Hsp90, the CK2 subunits were first deleted in SN95. To do so, both open reading frames of *CKA1*, *CKA2*, *CKB1*, and *CKB2* were deleted using amplicons of the nourseothricin recyclable marker encoded on plasmid pSD3 (pJK863[2]). Amplicons with 100 bp homology to the up- and down-stream regions of the respective ORF were generated using Q5 Taq Polymerase (Table S2) and transformed into SN95 as described above. Positive transformants were verified using genotyping PCR with the primers specified in Table S3. To epitope tag Hsp90 CK2 deletion strains YSD557 (*cka1*Δ/Δ), YSD623 (*cka2*Δ/Δ), YSD628 (*ckb1*Δ/Δ) and YSD634 (*ckb2*Δ/Δ), they were then transformed with the PCR amplified TAP-HIS1 construct off plasmid pSD18 (pFA-TAP-HIS)[3]. Successful TAP-tagging of Hsp90 was verified by diagnostic PCR using primers oSD86 and oSD522 (Table S3) and Western blotting.

### Supplementary Figures

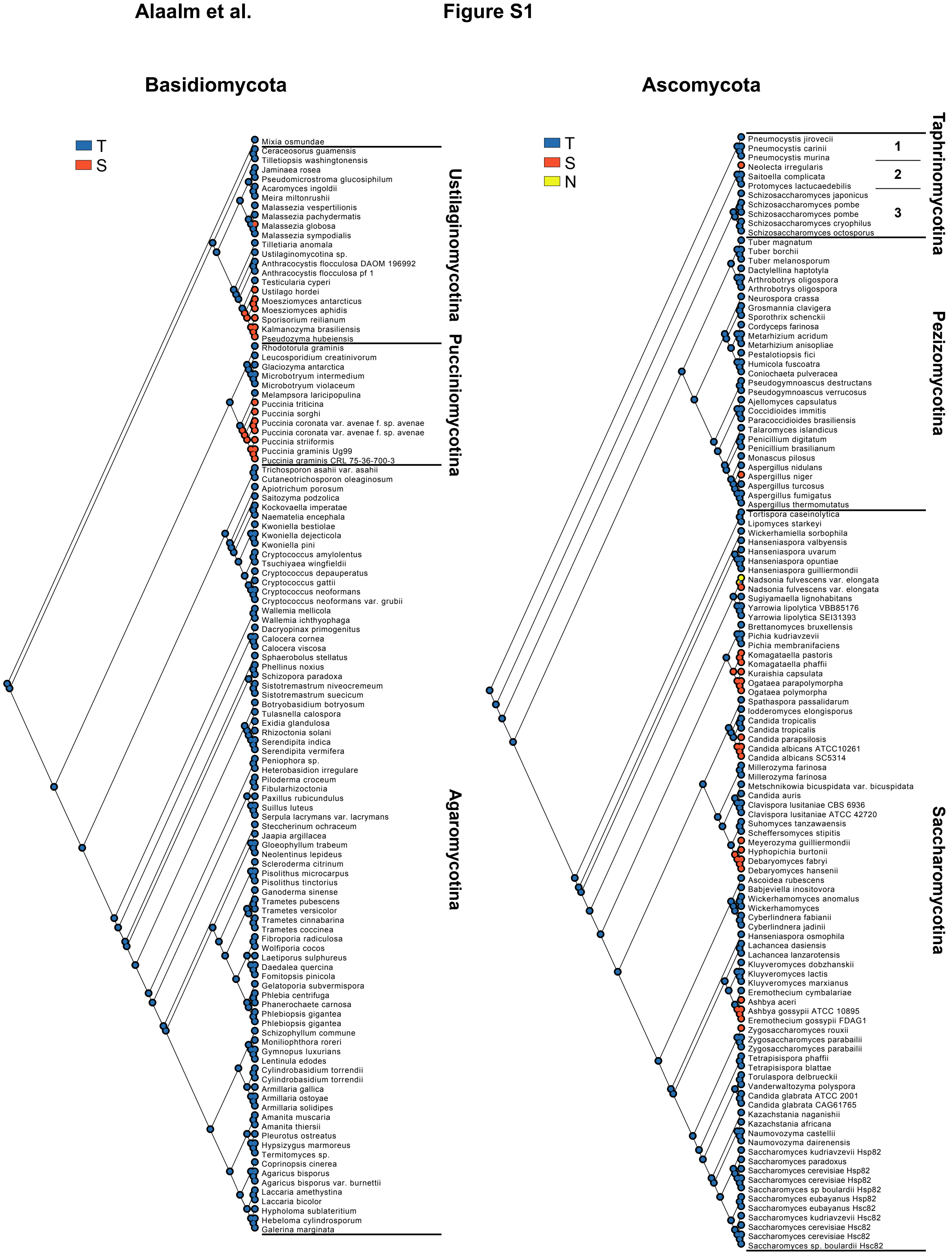


**Figure S1: The *HSP90*$^{S530}$ allele emerged multiple times independently during basidiomycota and ascomycota evolution.** Ancestral character state reconstruction using the Parsimony criterion for 116 basidiomycetous taxa and 119 ascomycetous taxa as depicted in the collapsed clades in Fig. 1c. Inset legends indicate colour coding used to distinguish between the different *HSP90* alleles. The *HSP90*$^{S530}$ allele has arisen ten-times independently in five of the six sub-phyla but could not be detected in the Agaromycotina.

### Figure S2

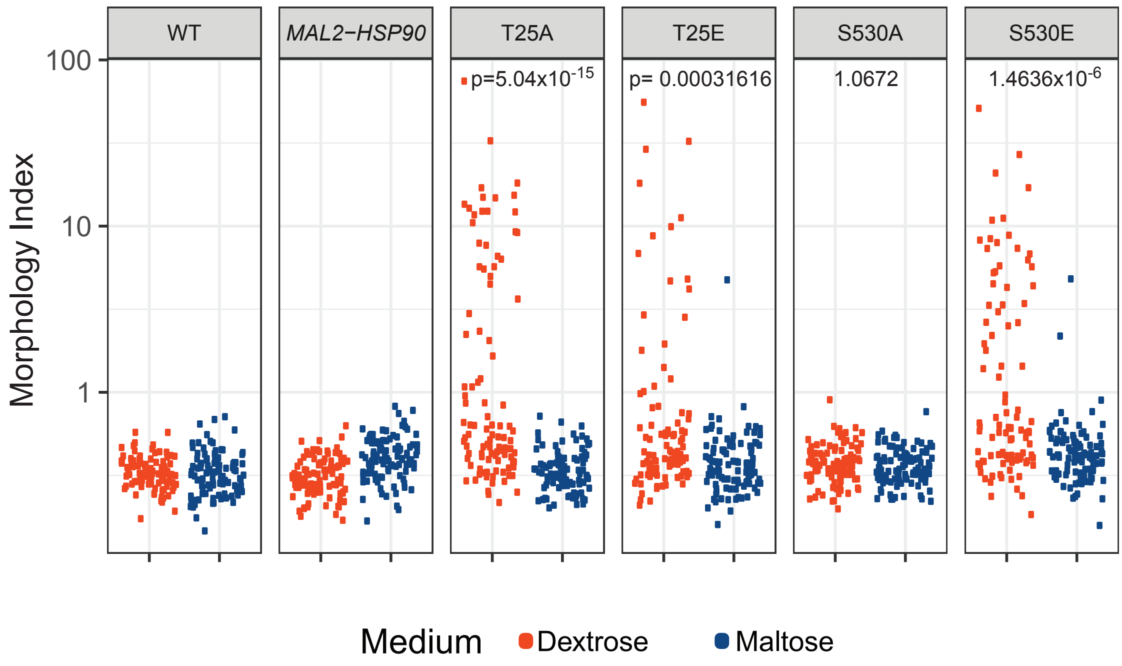


**Figure S2: Phosphorylation of S530 releases Hsp90's block on the yeast-to-hyphae transition.** *C. albicans* cells were grown in liquid rich media containing either maltose or dextrose and cell morphologies were analysed microscopically. 100 cells were selected randomly per strain and their morphology indices calculated. Morphology Indices, plotted on a logarithmic scale, were compared between cells expressing wild-type *HSP90* and those that express a mutant allele. Statistical significance was assessed using the Mann-Whitney U test. Alteration of the conserved T25 residue releases Hsp90's block on the yeast-to-hyphae transition. Cells of the phosphomimetic S530 are hyphal.
